# Intrinsic functional reorganisation of the attention network in the blind

**DOI:** 10.1101/245399

**Authors:** Wen Qin, Yong Liu, Tianzi Jiang, Chunshui Yu

**Author notes:** **Correspondence to:** Wen Qin, MD, Department of Radiology, Tianjin Medical University General Hospital, No. 154, Anshan Road, Heping District, Tianjin 300052, China.

## Abstract

Attention can bias visual perception by modulating the neuronal activity of visual areas. However, little is known if blindness can reshape the intrinsic functional organisation within the attention networks and between the attention and visual networks. A voxel-wise network-based functional connectivity strengthen mapping analysis was proposed to thirty congenitally, thirty early and thirty late blind subjects, and thirty sighted controls. Both the blind and sighted subjects exhibited similar spatial distributions of the intrinsic dorsal (DAN) and ventral (VAN) attention networks. Moreover, compared to the sighted controls, the blind subjects showed increased functional coupling within the DAN, and between the DAN and VAN, and between the attention sub-networks and visual areas, suggesting an increased information communication by visual deprivation. However, the onset age of blindness had little impact on the functional coupling of the attention network, indicating that non-visual sensory experience is enough for driving the development of intrinsic functional organization of the attention network. Finally, a positive correlation was identified between the duration of blindness and the functional coupling of the posterior inferior frontal gyrus with the visual network, representing an experience-dependent reorganisation after visual deprivation.

## Introduction

Attention is a cognitive process to focus our limited brain resources to preferentially process certain external stimuli or internal ideas while ignoring irrelevant ones (Raz and Buhle 2006). Attention is thought to be controlled by two segregated functional networks: a bilaterally distributed dorsal attention network (DAN) which is mainly composed of the frontal eye field (FEF) and intraparietal sulcus (IPS) and involves in endogenous and exogenous orienting of attention; and a right-lateralized ventral attention network (VAN) which is mainly consisting of the anterior insula (aINS) and temporoparietal junction (TPJ) and involves in reorienting of attention in response to salient stimuli (Corbetta and Shulman 2002; Corbetta et al. 2008). These two attention networks dynamically interact to determine which aspects of sensory information will be perceived (Buschman and Miller 2007; McMains and Kastner 2011). Both genetic and environmental factors can shape the organisation of the attention networks (Rueda et al. 2005; Bellgrove et al. 2007; Neufang et al. 2015). For example, visual attention training can improve the attention performance and reshape the attention networks in sighted subjects (Rueda et al. 2005; Rueda et al. 2012).

As blindness interrupts the main information transfer system of the human brain, long-term visual deprivation may affect attention behavior and reshape the organisation of the attention networks. In fact, superior auditory/tactile attention abilities have been observed in blind subjects (Roder et al. 1999; Collignon et al. 2006; Forster et al. 2007). The congenitally (CB) and early blind (EB) subjects have shown increased activation in the FEF and IPS of the DAN than the sighted control (SC) when performing attention-demanding tasks (Burton et al. 2004; Garg et al. 2007; Stevens et al. 2007; Burton et al. 2010). These findings indicate that visual deprivation can reshape regional activity of the attention networks. However, it is still unknown if and how visual deprivation reshapes the intrinsic functional couplings of the attention networks.

In the SC, the attention networks dynamically interact with the visual network (VN). Stimulus-driven signals from the VN can selectively recruit the VAN based on their salience and relevance; and top-down signals from the DAN can modulate the activity of visual areas to improve the perception of specific aspects of visual stimuli (Buchel et al. 1998a; Tootell et al. 1998; Jack et al. 2006; Silver et al. 2007; Cate et al. 2009; Bressler and Silver 2010). In the CB/EB, despite of the structural reorganisation (Shimony et al. 2006; Ptito et al. 2008; Jiang et al. 2009; Shu et al. 2009), the functional specification (location, motion, object, or language processing) of the occipital cortex is relatively preserved (Amedi et al. 2007; Renier et al. 2010; Collignon et al. 2011; Reich et al. 2011; Wolbers et al. 2011). However, it is unclear if the intrinsic functional organisation of the attention networks is also preserved in the blind. The cross-modal activation of the occipital cortex in the blind is frequently reported in the attention-demanding nonvisual tasks (Bavelier and Neville 2002; Collignon et al. 2009; Sathian and Stilla 2010), whereas simple passive auditory/tactile tasks cannot activate the occipital cortex (Sadato et al. 1996; Sadato et al. 1998; Weeks et al. 2000). Furthermore, the occipital cortex of the EB can be activated by pure top-down attention signals, which is very similar with the visual attention-induced response in the visual cortex in the SC (Gougoux et al. 2005; Stevens et al. 2007; Renier et al. 2010). These findings suggest that even after visual deprivation, functional interactions between the attentional and occipital regions still exist. However, little is known how visual deprivation impacts the intrinsic functional coupling between the attention and visual networks.

Visual deprivation-induced reorganisation may depend on the development maturity degree of the brain at the time of onset of blindness. Compared to the late blind (LB), the CB/EB exhibit different reorganisation patterns in cortical thickness (Jiang et al. 2009; Park et al. 2009; Kupers et al. 2011), glucose metabolism (Wanet-Defalque et al. 1988; Veraart et al. 1990), task-evoked activation (Buchel et al. 1998b), and functional connectivity density (FCD) (Qin et al. 2015) in the occipital cortex. Using the same auditory attention task, although both the EB and the LB demonstrated improvement in attention performance, the temporal and spatial alteration patterns in event-related potentials are largely different (Roder et al. 1999; Fieger et al. 2006). Nevertheless, we are unclear whether the developmental maturity degree of the brain at the time of onset of blindness has an effect on the intrinsic functional organisation of the attention networks in the blind.

Most of our knowledge on the functional organisation of the attention networks comes from the task-based functional magnetic resonance imaging (fMRI) that detects regional blood-oxygen-level-dependence (BOLD) activation evoked by a specific task (reflecting functional segregation). In contrast, the resting-state fMRI primarily focuses on the synchronization of spontaneous activity among brain regions (reflecting functional integration). A pioneer study has segregated the DAN and VAN by resting-state functional connectivity (rsFC) analysis (Fox et al. 2006). A following study further reveals a correlation between attention performance and the rsFC between the DAN and VAN in normal subjects (Wen et al. 2012). These findings indicate that rsFC analysis is a promising technique to evaluate the intrinsic functional organisation of the attention networks.

In this study, we recruited gender- and age-matched CB, EB, LB and SC (thirty subjects in each group), and proposed a voxel-wise network-based functional connectivity strength (FCS) mapping (NB_FCSM) method to compare intergroup FCS differences within each attention network (DAN and VAN), between the DAN and VAN, and between the attention and visual networks (DAN-VN and VAN-VN). Based on previous findings in the blind, we proposed four hypotheses: (1) the intrinsic functional organisation of the attention networks is preserved in the blind because the same brain regions show activation in non-visual attention tasks in the blind and in visual attention tasks in the SC (Garg et al. 2007; Stevens et al. 2007; Corbetta et al. 2008; Burton et al. 2010); (2) the blind subjects have increased functional couplings in the attention network since they show superior non-visual attention performance, and increased activation in the network during non-visual attention tasks (Roder et al. 1999; Collignon et al. 2006; Forster et al. 2007; Garg et al. 2007; Stevens et al. 2007); (3) the blind subjects have increased functional couplings between the attention and visual networks for they show increased occipital activation evoked by non-visual attention-demanding tasks (Gougoux et al. 2005; Stevens et al. 2007; Renier et al. 2010); and (4) the onset and/or duration of blindness may impact the intrinsic functional organisation of the attention networks in the blind as the CB/EB and LB show different reorganisation patterns in the occipital cortex (Jiang et al. 2009; Park et al. 2009; Collignon et al. 2013; Qin et al. 2015).

## Materials and methods

### Subjects

This study recruited 90 right-handed blind subjects and thirty sighted controls (22 males, age range 20-44 years). Blind subjects included thirty CB (onset since birth, 22 males, age range 18-39 years), thirty EB (onset from 0.5 to 12 years old, 22 males, age range 20-45 years), and thirty LB (onset from 13 to 28 years old, 22 males, age range 20-38 years). The samples size, age and gender of the four groups were well matched (Table 1). The protocol was approved by the Medical Research Ethics Committee of Tianjin Medical University, and written informed consent was obtained from all participants prior to the experiment.

**Table 1:** Demographic information of blind and sighted subjects

| Measures | CB | EB | LB | SC | Statistics | *P* value |
| --- | --- | --- | --- | --- | --- | --- |
| Gender (male/female) | 22 / 8 | 22 / 8 | 22 / 8 | 22 / 8 | $\chi^2 = 0^a$ | 1 |
| Age (mean ± SD), years | 25.7 ± 4.6 | 28.5 ± 6.6 | 28.2 ± 4.1 | 28.6 ± 5.9 | $F = 1.95$ | 0.125 |
| Onset age of blindness (mean ± SD), years | 0 | 7.7 ± 3.2 | 18.0 ± 4.4 | / | / | / |
| Duration of blindness (mean ± SD), years | 25.7 ± 4.6 | 20.8 ± 7.9 | 10.2 ± 4.2 | / | / | / |
| Mean FD (mean ± SD) | 0.15 ± 0.07 | 0.16 ± 0.05 | 0.15 ± 0.07 | 0.14 ± 0.05 | $F = 0.21$ | 0.888 |
| Motion spikes (mean ± SD), % | 1.69 ± 2.54 | 1.59 ± 2.35 | 2.31 ± 4.35 | 0.94 ± 1.79 | $\chi^2 = 2.33^b$ | 0.508 |

Note: <sup>a</sup> chi-square test. <sup>b</sup> Kruskal Wallis Test, A motion spike is defined as a time point with FD > 0.5. CB = congenitally blind, EB = early blind, FD = frame-wise displacement, SC = sighted control, SD = standard deviation.

### MRI data acquisition

MRI data were obtained using a 3.0 Tesla MR scanner (Trio Tim system; Siemens, Erlangen, Germany) with a 12-channel head coil. The resting-state fMRI data were acquired using a gradient-echo echo-planar imaging sequence with the following parameters: repetition time = 2000 ms, echo time = 30 ms, flip angle = 90°, matrix = 64 × 64, field of view = 220 mm × 220 mm, 32 axial slices with a thickness of 3 mm and a gap of 1 mm. During scans, all subjects were instructed to keep eyes closed and awake, stay motionless, and think of nothing in particular. The three-dimensional structural images were acquired using a magnetization-prepared rapid-acquisition gradient echo sequence: repetition time = 2000 ms, echo time = 2.6 ms, inversion time = 900 ms, flip angle = 9°, matrix = 256 × 224, field of view = 256 mm × 224 mm, 176 continuous sagittal slices with a thickness of 1 mm.

### The fMRI data preprocessing

The resting-state fMRI data were preprocessed using Statistical Parametric Mapping (SPM8, http://www.fil.ion.ucl.ac.uk/spm). The first 10 volumes were removed because of incomplete T1 relaxation, and to allow the participants to adapt to the scanning environment. The remaining 170 volumes were corrected for the acquisition time delay between slices and were realigned to the first volume to estimate and correct rigid head motion. All subjects were within the predefined head motion thresholds of translation < 2 mm and rotation < 2°. We also calculated the frame-wise displacement (FD) (Power et al. 2012) and percent of spike motion (defined as FD > 0.5) in each subject. The mean FD was included as an additional nuisance variable during all fMRI-related statistical analyses to further exclude out the effect of head motion. Then fMRI dataset were affinely corregistered with the structural volume, and the structural images were segmented and nonlinearly normalized into the Montreal Neurological Institute (MNI) space. Finally, the fMRI volumes were written to the MNI space and resampled into 3-mm^3^ voxels using the deformation parameters derived from structural images. Several spurious variances, including the motion parameters (six parameters and their first time derivatives), and average BOLD signals of the whole brain, ventricular and white matter were removed from the normalized fMRI data through linear regression. After band-pass frequency filtering (0.01-0.08 Hz), the functional images were spatially smoothed with a Gaussian kernel of 6 mm full width at half maximum.

### Attention network extraction

The attention networks were extracted based on a meta-analysis of attention-evoked coactivation. BrainMap Sleuth 2.0.3 (http://www.brainmap.org/) was used to search papers reporting brain activation evoked by visual attention tasks in right-handed healthy adults using either positron emission tomography or fMRI. We excluded conditions aimed to investigate the effects of diseases, handedness, gender or drugs on task-evoked activation. All retrieved foci were transformed into MNI space and manually checked to exclude those out of the MNI grey matter mask. A total of 128 papers that included 360 experiments, 4793 subjects, and 3470 foci were finally included in the meta-analysis.

The meta-analysis was performed using the revised version of Activation Likelihood Estimation (ALE) technique (Eickhoff et al. 2009; Turkeltaub et al. 2012) implemented in GingerALE 2.3.6 (http://www.brainmap.org/). This algorithm aims to identify regions showing convergent activation across experiments if the merged activation is higher than the expectation under a random spatial association. For a given experiment, all activation coordinates are modelled as independent Gaussian probability distributions and combined to generate a modelled activation (MA) map for this experiment. To minimize within-experiment foci number and proximity on the ALE calculation, the maximum probability of each voxel in a given experiment was considered as the MA value (Turkeltaub et al. 2012). Then, the ALE score of each voxel was calculated by summing individual MA values of all experiments. A random-effect non-parametric permutation test was used to identify voxels with significant ALE differences against the null-hypothesis. Multiple comparisons were corrected using a false discovery rate (FDR) method (*q* < 0.05 and cluster size > 1000 mm^3^). The task-evoked spatial activation pattern in the attention network was shown in Supplementary Fig. 1.

### DAN and VAN extraction

As performed in a prior study (Fox et al. 2006), we used rsFC-based conjunction analysis to define the DAN and VAN. The right FEF [28 −6 52] and IPS [30 −58 46] were used to identify the DAN, and the right aINS [38 26 −6] and TPJ [58 −42 30] were used to identify the VAN. These peak MNI coordinates were defined based on the ALE map (Supplementary Fig. 1). A 6mm-radius sphere centered on each coordinate was drawn and the overlap voxels between the sphere and ALE map were taken as the seed. Pearson correlation coefficients between the mean time series of each seed and that of each voxel of the whole brain were computed and transformed into *z*-values using Fisher’s transform. In each group, individuals’ z-values were entered into a one-sample *t*-test to identify brain regions with significant positive correlations with each seed. The conjunction analysis was then applied to identify the DAN regions that exhibit positive rsFC with both the FEF and IPS and the VAN regions that show positive rsFC with both the aINS and TPJ (*q* < 0.05, FDR corrected; cluster size > 30 voxels). The spatial distributions of the DAN and VAN of each group are shown in Supplementary Figs 2 and 3, respectively. We used a leave-one-out (LOO) method to verify if the inter-subject variability would impact the spatial distribution pattern of the DAN and VAN of each group. In each LOO experiment, we performed conjunction analysis using the same method as the original one after excluding one subject’s rsFC data. The spatial overlapping ratio (SOR) and spatial correlation coefficient (SCC) of the DAN/VAN between each LOO experiment and original one were calculated. The SOR and SCC of the DAN/VAN between each pair of groups were also calculated to evaluate the spatial similarity of the attention networks among the groups.

To increase the functional specificity of the DAN and VAN, we further redefined the two attention networks within brain regions that showed both high functional connectivity with the hubs of this network and high activation by visual attention tasks, which was computed by multiplying the rsFC-based DAN and VAN with the ALE-based coactivation map. Because we aimed to investigate if the intrinsic functional couplings of brain regions that originally serve visual attention in the SC would change after visual deprivation, the redefined DAN and VAN were only calculated from the SC subjects.

### NB-FCSM

A NB-FCSM method was proposed to voxel-wisely compare intergroup FCS differences within (DAN and VAN) and between networks (DAN-VAN, DAN-VN and VAN-VN). Similar to the FCD mapping that measures the degree distribution of the intrinsic functional networks at the voxel-level (Tomasi and Volkow 2012), NB_FCSM measures the weighted degree distribution, namely the FCS (Liang et al. 2013). As an extension of the original whole-brain FCS analysis (Liang et al. 2013), this method can not only measure FCS distribution within a network, but also map the FCS between each pair of networks, which can provide information about inter-network functional coupling. The preprocessing steps were the same as the rsFC analysis. The pipeline of NB-FCSM is shown in Fig. 1. For within-network FCS, we first calculated the rsFC between each pair of voxels within a given network to construct a voxel-wise correlation matrix. Then a threshold (*P* < 0.05, uncorrected) of the positive rsFC was used to sparsify the correlation matrix. Within-network FCS of a given voxel was calculated as the average rsFC between this voxel and all other survived voxels within the sparse matrix, and this step was iterated for each remaining voxel. Similarly, between-network FCS was calculated except that the rsFC was calculated from each pair of voxels from two different networks (e.g., A and B). We obtained two between-network FCS measures: FCS_A&B_ represents the average rsFC of one voxel in network A with all survived voxels in network B, and vice versa for FCS_B&A_. To test the effect of connectivity thresholding on our results, we also calculated FCS using four different thresholds (r > 0, 0.1, 0.2 and 0.3) and compared the intergroup differences.

**Figure 1.**
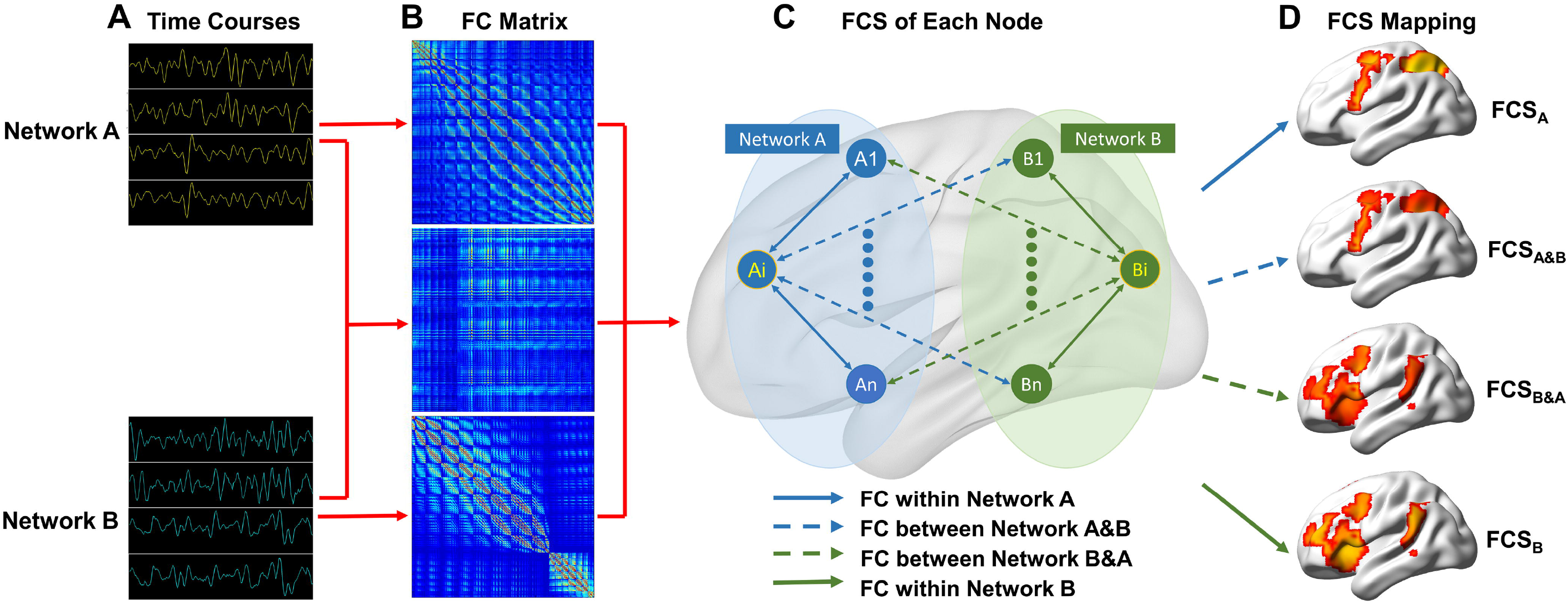
Pipeline of network-based functional connectivity mapping. Abbreviations: FC = functional connectivity, FCS = functional connectivity strength. There are four main steps for NB-FCSM: A) Extracting the timecourses of each voxel for each network; B) calculating the functional correlation matrix within each network and between each pair of networks; C) calculating the functional connectivity strength of each voxel; D) generating the FCS maps of each network and between each pair of networks.

### Statistical analyses

A fixed-effect general linear model (GLM) was used to voxel-wisely compare intergroup differences in FCS within each attention network, between the DAN and VAN, and between the DAN/VAN and VN (*q* < 0.05, FDR corrected), controlling for the effects of age, gender and mean FD. Brain regions with significant FCS differences were extracted as regions-of-interest (ROIs) and entered into *post-hoc* comparisons to test pair-wise differences among the four groups (*q* < 0.05, FDR corrected). To clarify which functional connectivities contributed to the FCS changes in blind subjects, the rsFC between each identified ROI and each voxels of the whole brain were calculated, and intergroup differences in rsFC were compared using the same statistical model as the FCS analyses (*q* < 0.05, FDR corrected). Finally, partial correlation analyses controlling for the effects of gender and mean FD were performed to test possible correlations of regional FCS with onset age and duration of blindness (*q* < 0.05, FDR corrected).

## Results

### Demographic data

The demographic data of these subjects are shown in Table 1. A total of thirty CB, thirty EB, thirty LB and thirty SC were included in this study. There were no significant differences in gender (χ^2^ = 0, *P* = 1), age (*F* = 1.95, *P* = 0.125), mean FD (*F* = 0.21, *P* = 0.888) and percentage of motion spikes (χ^2^ = 2.33, *P* = 0.508) among the four groups. The age of onset of blindness was 0 year in the CB, 7.7 ± 3.2 years in the EB, and 18.0 ± 4.4 years in the LB; and the duration of blindness was 25.7 ± 4.6 years in the CB, 20.8 ± 7.9 years in the EB, and 10.2 ± 4.2 years in the LB.

### The spatial distribution of the attention network

The spatial distribution of the attention network identified by coordinated-based ALE meta-analysis (*q* < 0.05, FDR corrected) is shown in Supplementary Fig. 1. Visual attention-evoked activation was located in the bilateral FEF, IPS, middle (MFG) and inferior (IFG) frontal gyrus, superior parietal lobule (SPL), dorsal anterior cingulate cortex (dACC), aINS, and the right TPJ. The spatial distributions of the DAN and VAN in each group identified by rsFC-based conjunction analyses (*q* < 0.05, FDR corrected) are shown in Fig. 2A-C, and Supplementary Figs. 2 and 3. The spatial distribution maps of these two networks were similar among the CB, EB, LB and SC (SOR: 56.3 ± 4.7% for the DAN and 55.2 ± 2.2% for the VAN; SCC: 0.693 ± 0.038 for the DAN and 0.524 ± 0.048 for the VAN). The overlapped regions of the DAN for all groups included the bilateral FEF, IFG, IPS and SPL, whereas the overlapped regions of the VAN included the right-lateralized TPJ, aINS, IFG, MFG, dACC and superior frontal gyrus (SFG). LOO methods demonstrated that the spatial distributions of the rsFC-based DAN/VAN were highly consistent in each group (SOR = 91.7 ± 1.6%, SCC = 0.991 ± 0.004), indicating a highly reproducibility of the resting-state DAN and VAN (Fig. 2D). The spatial distributions of the redefined DAN and VAN with both positive rsFC and high activation probability in the SC are shown in Fig. 3. The DAN was composed of the bilateral FEF, posterior IFG (pIFG), IPS and SPL; and the VAN was a right-lateralized network and included the aINS, TPJ, pIFG, MFG, and dACC. The right pIFG was the only region shared by the DAN and VAN.

**Figure 2.**
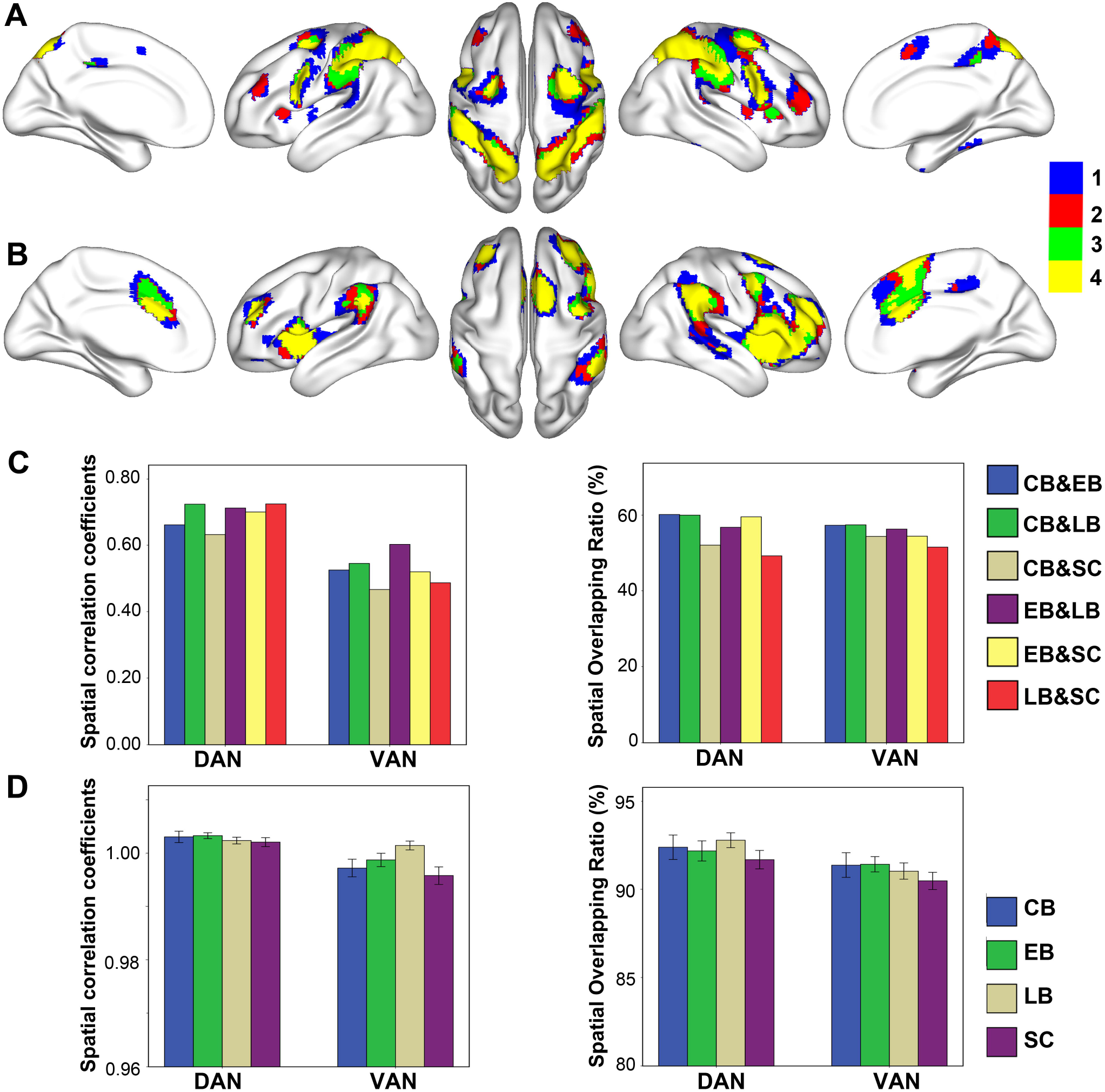
Spatial similarities of the dorsal and ventral attention networks derived from rsFC-based conjunction analysis. **(A)** and **(B)** show spatial overlaps of the DAN and VAN across groups, respectively. The color of each voxel indicates the number of groups (1 to 4) that share this voxel in the DAN or VAN. **(C)** represents the spatial correlation coefficients and overlapping ratio of the attention networks between each pair of groups. **(D)** shows the spatial correlation coefficients and overlapping ratio of the attention networks between each LOO iteration and the use of full data in each group.

**Figure 3.**
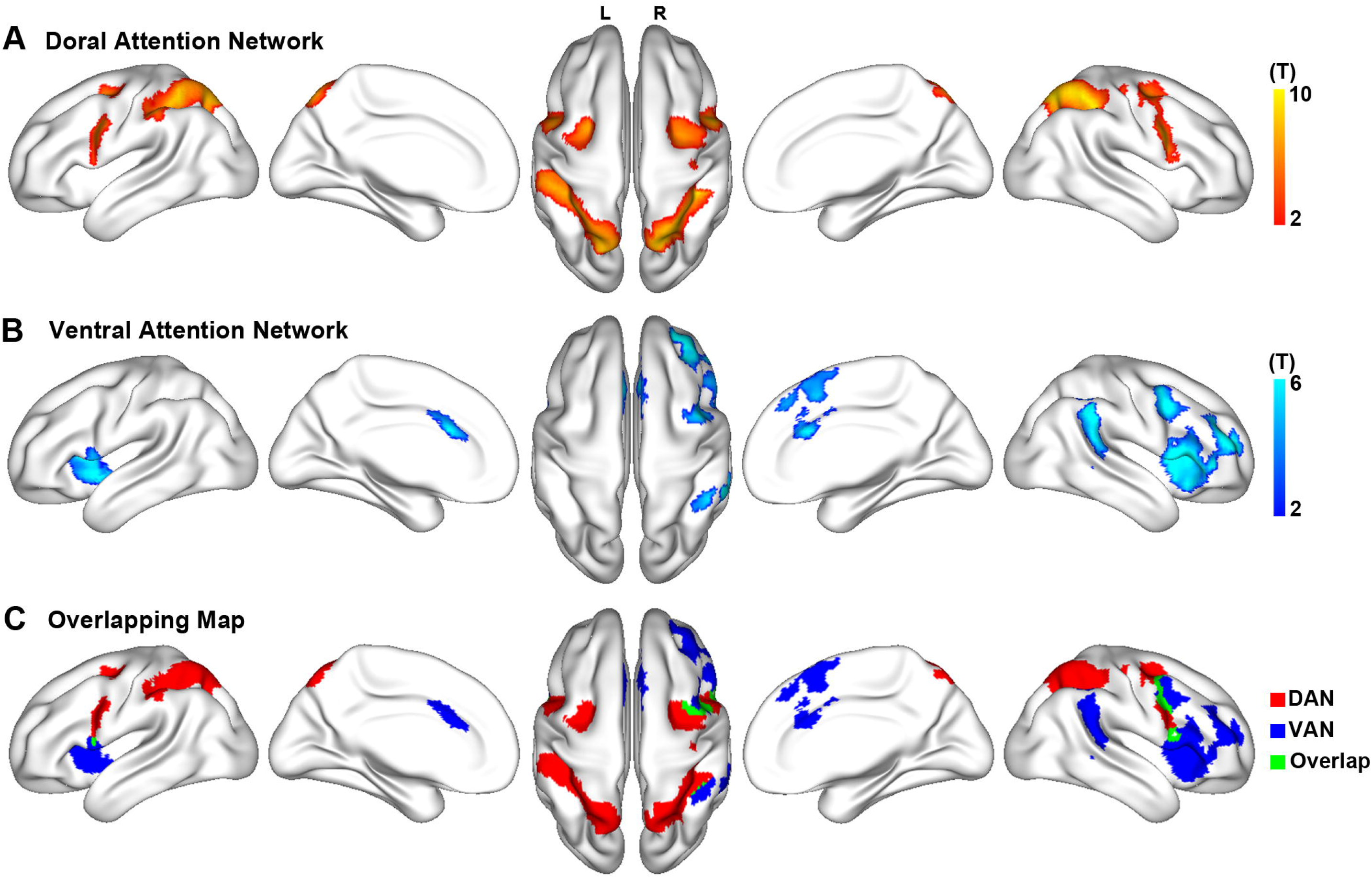
The refined dorsal and ventral attention networks in the sighted controls. The DAN **(A)** and VAN **(B)** were identified by combination of ALE-based coactivation and rsFC-based conjunction analyses (*q* < 0.05, FDR corrected). Each non-zero voxel of an attention network has both positive rsFC with the hubs of this network and high activation probability by visual attention tasks. The color bar represents the T value of the conjunction analysis. The DAN (red) and VAN (blue) are spatially overlapped at the right posterior IFG (green) **(C).**

### FCS changes within the attention network in the blind

Within the DAN, GLM analysis showed significant intergroup FCS differences (*q* < 0.05, FDR corrected) in the bilateral SPL/IPS and the right FEF and pIFG (Fig. 4A). The blind groups generally had increased FCS than the SC group, but they did not differ from each other. There were no significant FCS differences within the VAN among the four groups.

**Figure 4.**
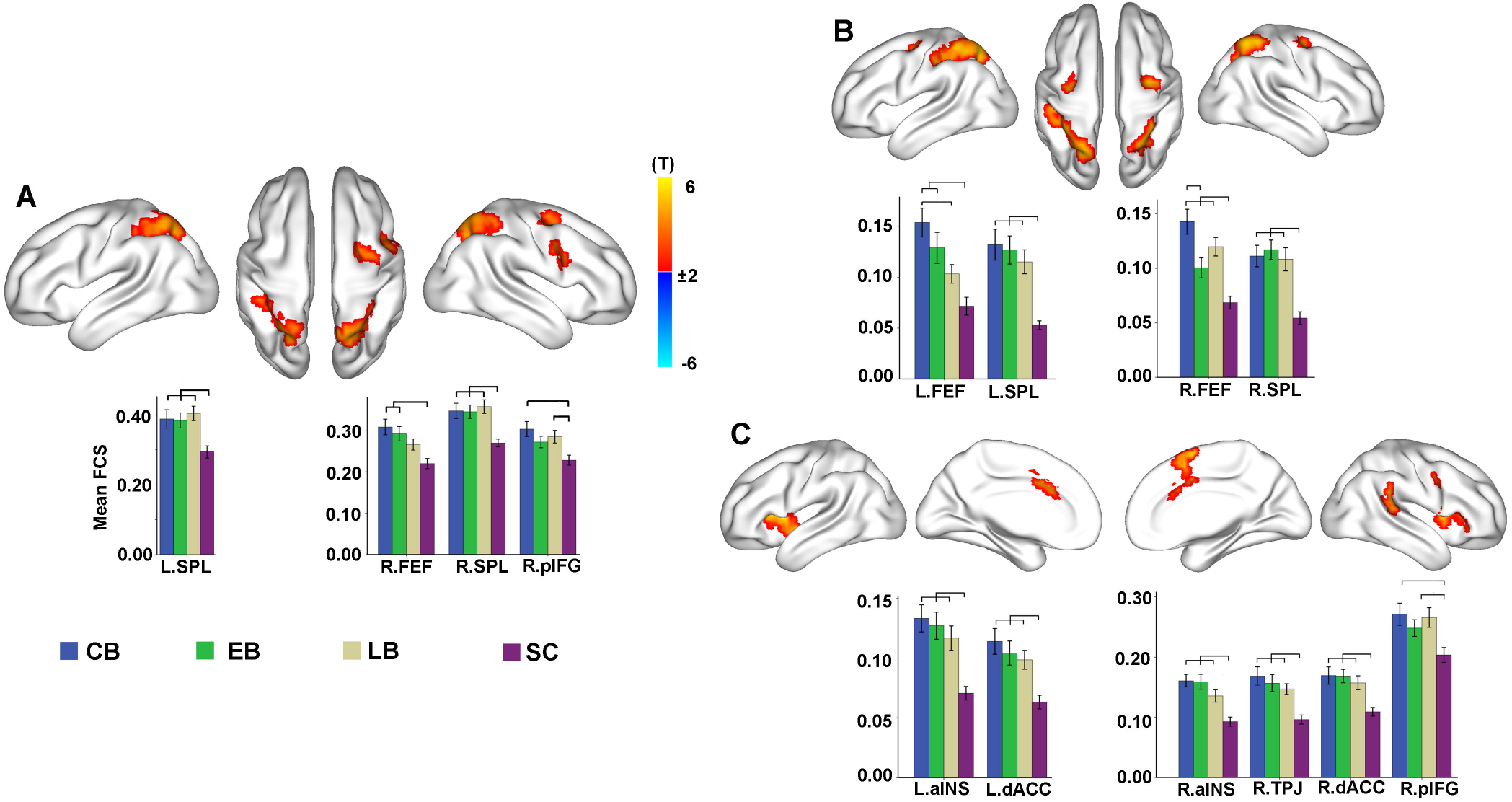
FCS changes within the attention networks in the blind. General linear model is used to compare FCS differences within the attention networks between the total blind subjects and the sighted controls (*q* < 0.05, FDR corrected), while controlling for the effects of age, gender, and mean FD. Color bar represents the T value. Warm and cold colors represent increased and decreased FCS in the blind, respectively. **(A)** shows the intergroup FCS differences within the DAN; **(B)** shows brain regions of the DAN that exhibit intergroup differences in FCS with the VAN; and **(C)** shows brain regions of the VAN that exhibit intergroup differences in FCS with the DAN. The lines connecting any two groups in bar graphs indicate significant differences between the two groups (*q* < 0.05, FDR corrected).

Significant between-network FCS differences (*q* < 0.05, FDR corrected) were found between the bilateral FEF and SPL/IPS (DAN components) and the VAN (Fig. 3B), and between the bilateral aINS, dACC, the right pIFG and TPJ (VAN components) and the DAN (Fig. 3C). Generally, the FCS in each region was increased in the blind than in the SC, but there were no significant differences among the blind groups. The only exception was that the CB had a higher FCS between the right FEF and the VAN than the EB, and a higher FCS between the left FEF and the VAN than the LB (*q* < 0.05, FDR corrected).

In the validation analysis, we recalculated FCS using four different connectivity thresholds (r > 0, 0.1, 0.2 and 0.3, respectively) and repeated these intergroup comparisons. The intergroup FCS differences were highly consistent across the four thresholds and very similar with the original ones (Supplementary Figs 4 and 5).

### FCS changes between the attention and visual networks in the blind

The GLM showed significant intergroup FCS differences (*q* < 0.05, FDR corrected) between the DAN and VN (Fig. 5A and B). Generally speaking, compared to the SC, the blind subjects showed increased FCS between hubs (the bilateral FEF, IPL and pIFG) of the DAN and the VN and between higher-level visual hubs (the bilateral superior [SOG] and inferior [IOG] occipital gyri, the left lingual gyrus [LG] and the right middle occipital gyrus [MOG]) and the DAN, and decreased FCS between the primary visual areas (the bilateral calcarine sulcus [CalS]) and the DAN (most of them can pass FDR correction). Among the blind subjects, the CB had a lower FCS between the left IOG and the DAN compared to the EB and LB, while higher FCS between the right pIFG and the VN relative to the LB (*q* < 0.05, FDR corrected) (Fig. 5A and B).

**Figure 5.**
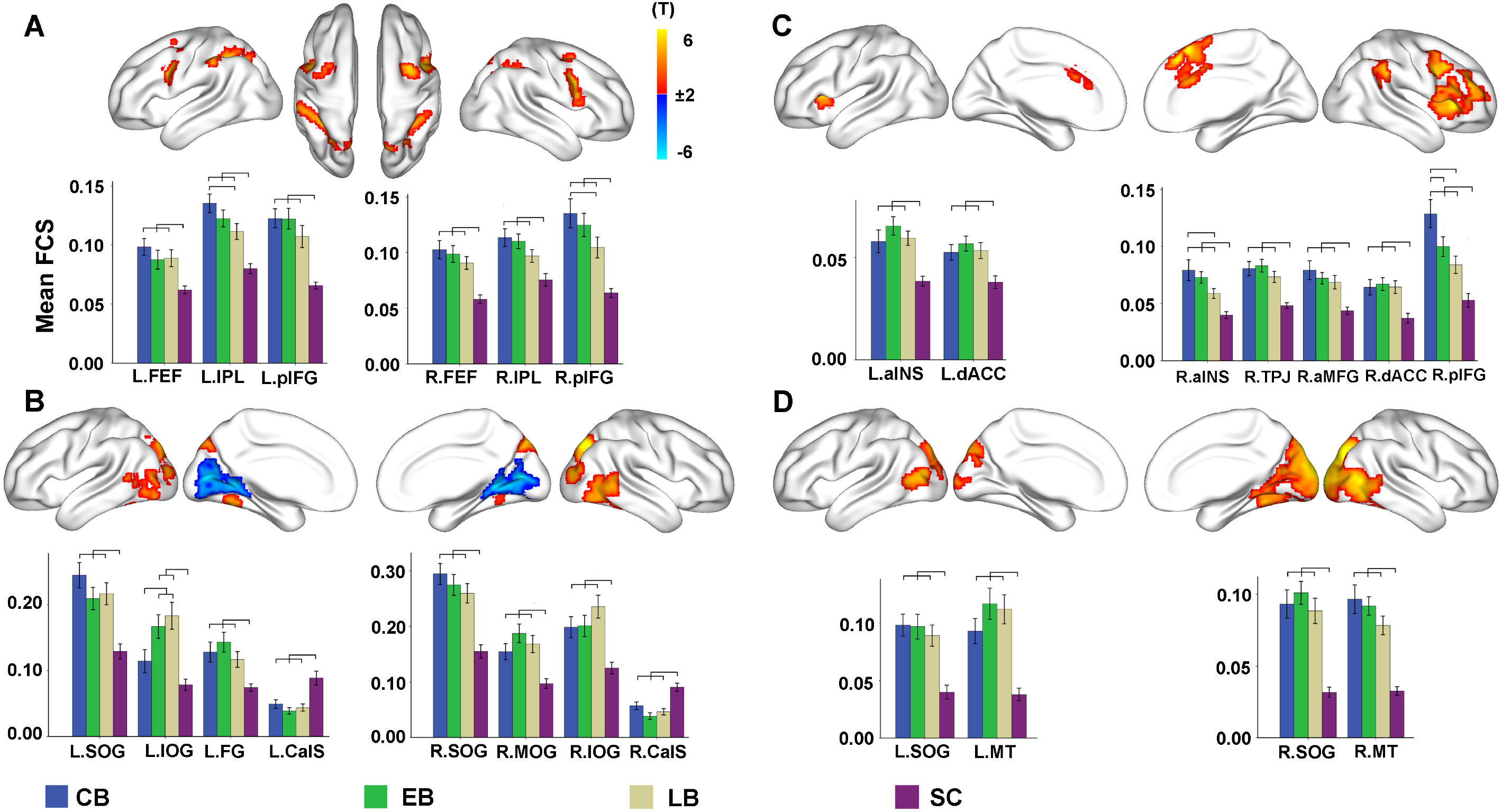
FCS changes between the attention and visual networks in the blind. General linear model is used to compare FCS (between the attention and visual networks) differences between the total blind subjects and the sighted controls (*q* < 0.05, FDR corrected), while controlling for the effects of age, gender, and mean FD. Color bar represents the T value. Warm and cold colors represent increased and decreased FCS in the blind, respectively. **(A)** shows brain regions of the DAN that exhibit intergroup differences in FCS with the VN; **(B)** shows brain regions of the VN that exhibit intergroup differences in FCS with the DAN; **(C)** shows brain regions of the VAN that exhibit intergroup differences in FCS with the VN; **(D)** shows brain regions of the VN that exhibit intergroup differences in FCS with the VAN. The lines connecting any two groups in bar graphs indicate significant differences between the two groups (*q* < 0.05, FDR corrected).

GLM analysis also revealed significant intergroup FCS differences (*q* < 0.05, FDR corrected) between the VAN and VN (Fig. 5C and D). Compared to the SC, the blind subjects showed increased FCS between hubs (the right pIFG, MFG, TPJ, and the bilateral aINS and dACC) of the VAN and the VN and between hubs (most of the right visual areas, the left middle temporal visual area [MT] and SOG) of the VN and the VAN (*q* < 0.05, FDR corrected). No significantly decreased FCS between the VAN and VN were found in any blind groups. In the blind groups, the CB showed a higher FCS between the right aINS and the VN than the LB (*q* < 0.05, FDR corrected), and a higher FCS between the right pIFG and the VN than the EB and LB (*q* < 0.05, FDR corrected) (Fig. 5C and D). As shown in Supplementary Figs 4-6, the intergroup FCS differences between the attention and visual networks were highly consistent across rsFC thresholds and very similar with the original ones.

### The rsFC changes within the attention network in the blind

Treating all regions with significant FCS differences as seeds, voxel-wise rsFC analyses (*q* < 0.05, FDR corrected) demonstrated: within the DAN, the blind subjects generally had increased rsFC between the left SPL and the left IPS and pIFG, and between the right SPL and the right FEF, pIFG and IPS compared to the SC. Blind subjects also had increased rsFC between most hubs (such as the bilateral FEF and SPL) of the DAN and those (such as the bilateral aINS and dACC, and right pIFG and TPJ) of the VAN. However, there were no significant differences in rsFC within the attention network among blind subjects (Supplementary Fig. S7).

### The rsFC changes between the attention and visual networks in the blind

Voxel-wise rsFC analyses (*q* < 0.05, FDR corrected) demonstrated that most hubs (including bilateral FEF, IPL and pIFG) of the DAN had increased rsFC with higher-level visual areas in the blind compared with the SC, while the primary visual areas (bilateral CalS) showed decreased rsFC with most DAN hubs (Supplementary Fig. S8). Blind subjects also had increased rsFC between most hubs (such as the bilateral dACC and aINS, and right MFG, pIFG, and TPJ) of the VAN and those (such as the bilateral MT and SOG) of the VN (Supplementary Fig. S9). There were no significant differences in rsFC of DAN-VN and VAN-VN among blind subjects.

### Correlations between connectivity changes and blindness chronometry

While controlling for gender and head motion effects, partial correlation analyses (*q* < 0.05, FDR corrected) showed that duration of blindness was positively correlated with the FCS between the right pIFG of both the VAN (*pr* = 0.346, *P* < 0.001) and DAN (*pr* = 0.311, *P* = 0.003) and the VN, and negatively correlated with the FCS between the left IOG of the VN and the DAN (*pr* = −0.326, *P* = 0.002) (Fig. 9). Furthermore, after additionally controlling for the onset of blindness, the correlations between the right pIFG of the VAN and the VN and between the left IOG of the VN and the DAN were still significant. There were no significant correlations between the FCS of each ROI and the age of onset of blindness (*q* < 0.05, FDR corrected).

**Figure 6.**
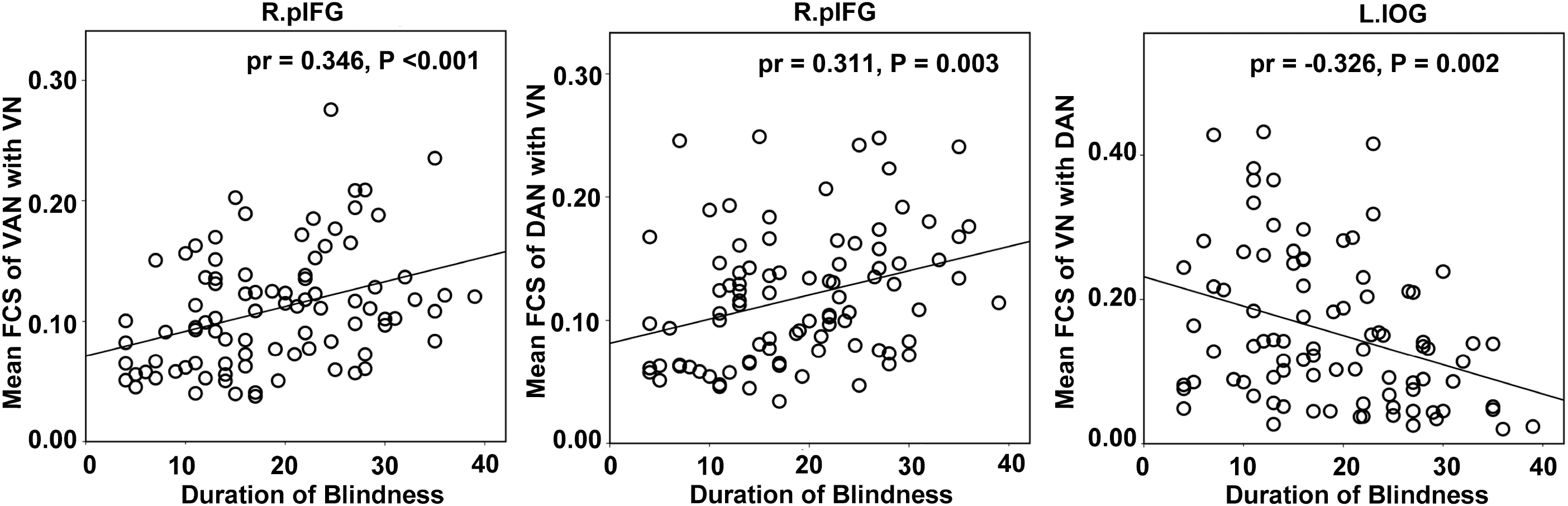
Correlations between duration of blindness and FCS in the blind. Partial correlation analyses controlling for the effects of gender and mean FD are performed to test correlations between duration of blindness and regional FCS in the blind (*q* < 0.05, FDR corrected).

## Discussion

In this study, we found that the blind had similar spatial pattern of the attention network with the SC, suggesting that the intrinsic functional organisation of the attention network is preserved in the blind. The blind subjects showed increased functional connectivity within the DAN, between the DAN and VAN, and between the attention and visual networks, indicating increased information communication within and between these attention-related networks in the blind. The lack of connectivity differences between the blind groups and the lack of correlations between connectivity and onset age of blindness suggest that the intrinsic functional organisation of the attention network is not influenced by its developmental maturity degree at the time of onset of blindness. Our study provided systematic knowledge of the intrinsic functional reorganisation of the attention networks by blindness with different onset age, and can improved our understanding about the interactions between higher-level attention network and visual areas in compensating for visual loss..

### Methodological consideration

In light of the FCD that voxel-wisely measure the unweighted degree distribution of the whole brain network (Tomasi and Volkow 2012), the FCS is proposed to voxel-wisely measure the weighted degree distribution of the network (Liang et al. 2013). In contrast to the FCD that is sensitive to connection thresholds, the FCS is insensitive to connection thresholds because of the weighting property, which is supported by our findings of different connection thresholds resulting in similar intergroup FCS differences. Here, we extended the traditional whole-brain-level FCS analysis to voxel-wisely assess the FCS within and between networks, namely the NB-FCSM. Compared with the independent component analysis (ICA) that is typically used to measure the global between-network functional connectivity (Allen et al. 2014; Wang et al. 2014), our voxel-wise between-network FCS analysis can identify the specific brain regions with between-network connectivity differences between groups. A prerequisite for robust application of NB-FCSM is accurate definition of networks of interest. Here, we combined the coordinated-based ALE meta-analysis and the rsFC-based conjunction analysis to more accurately define the DAN and VAN. Thus, each voxel of the redefined attention networks possess both high functional connectivity and coactivation, which was used for NB-FCSM.

### Preserved intrinsic functional organisation in the attention network in the blind

The human attention system closely interacts with the sensory systems. For example, the VAN controls stimulus-driven reorienting from sensory modalities and the DAN generates top-down signals to bias the response of the sensory cortex (Astafiev et al. 2003; Kincade et al. 2005) (Astafiev et al. 2006; Indovina and Macaluso 2007). As the main information input of the human brain, visual experience plays an important role in establishing and reshaping the functional organisation of the attention network. For example, visual attention training can reshape the attention network in normal subjects (Rueda et al., 2005; Rueda et al., 2012). In this study, we found that the attention networks showed similar spatial distribution in the blind and the SC, suggesting that the intrinsic functional organisation of the attention networks is preserved in the blind. This finding also indicates that the functional organisation of the attention network can be normally established in the blind by receiving inputs from non-visual sensory modalities because there are dense connections between the attentional regions and the thalamus (Behrens et al. 2003) and non-visual sensory cortices (Umarova et al. 2010). The preserved functional organisation of the attention network in the blind may provide new evidence for the supramodal nature of the attention regions, which is supported by the sensory modality-independent functional specialization of the attention network. For example, a certain attentional region is functionally specialized for processing a specific type of attention despite of the inputs from what sensory modalities (Downar et al. 2000; Smith et al. 2010; Ptak 2012).

### Enhanced intrinsic functional coupling of the attention network in the blind

Superior auditory/tactile attention performance has been frequently reported in the blind (Roder et al. 1999; Collignon et al. 2006; Forster et al. 2007; Collignon and De Volder 2009). Enhanced activation in the FEF and IPS is also observed in the CB/EB compared the SC when performing attention-demanding tasks (Burton et al. 2004; Garg et al. 2007; Stevens et al. 2007; Burton et al. 2010). In this study, we found increased functional couplings within and between the attention networks in the blind, which may provide an explanation for the superior attention performance from a new perspective of functional integration. We found increased connectivity in the DAN but not in the VAN in the blind, which is consistent with previous studies reporting enhanced activation only in the DAN regions in the CB/EB than the SC during attention-demanding tasks (Burton et al. 2004; Garg et al. 2007; Stevens et al. 2007; Burton et al. 2010). This finding suggests that the DAN and VAN respond differently to visual deprivation, reflecting different dependencies of the development of the functional organisation of the DAN and VAN on visual experience.

The DAN and VAN do not work independently, instead, they closely interact to determine which aspects of sensory information will be attended to (Buschman and Miller 2007; McMains and Kastner 2011). In consistent with a previous study (Fox et al. 2006), we found that the right posterior IFG/MFG was the region shared by the DAN and VAN, which may be the anatomical substrate for information communication between the two attention networks. In the theory of reorienting (Corbetta et al. 2008), when one focuses on an object, the DAN sends sustained top-down inhibition signals to the right TPJ to prevent unimportant and low-relevant stimuli from transferring to the VAN; when the stimuli are important and relevant enough to break the inhibition by the TPJ, the VAN is activated and sends the reorienting salient signals to the DAN. The increased intrinsic functional coupling between the DAN and VAN suggests that the information transfer efficiency or the functional integration of the attention networks is enhanced after visual deprivation, which may facilitate the switching between top-down attention and stimulus-driven reorienting. This finding may explain why the blind subjects have superior attention performance (Roder et al. 1999; Collignon et al. 2006; Forster et al. 2007; Collignon and De Volder 2009).

### Enhanced intrinsic functional coupling between the attention and visual networks in the blind

In the SC, the DAN strongly interacts with the visual areas to bias the visual perception (Buchel et al. 1998a; Tootell et al. 1998; Jack et al. 2006; Silver et al. 2007; Cate et al. 2009; Bressler and Silver 2010). In the blind, the deprived visual perception of the V1 may cause reduced functional coupling between the DAN and the V1. In contrast, the V1 of the blind involves in attention-demanding nonvisual tasks (Bavelier and Neville 2002; Collignon et al. 2009; Sathian and Stilla 2010), which may cause an increased functional coupling between the DAN and the V1. Thus, the reduced functional coupling between the DAN and the V1 in the blind may be a combined consequence of these two mechanisms. The same mechanisms can also be applied to explain for the increased functional coupling between the VAN and the V1 in the blind. In contrast to the V1, the functional specialization of the higher-level visual areas has found to be preserved to process non-visual stimuli in the blind (Amedi et al. 2007; Renier et al. 2010; Collignon et al. 2011; Reich et al. 2011; Wolbers et al. 2011). They can interact with the attention networks to process non-visual attention tasks after visual deprivation (Buchel et al. 1998a; Tootell et al. 1998; Jack et al. 2006; Silver et al. 2007; Cate et al. 2009; Bressler and Silver 2010). Thus, our findings of the increased functional coupling between the attention and higher-level visual areas in the blind may reflect an increased interaction between them, which may provide an explanation for the increased attention task-evoked activation (Garg et al. 2007; Stevens et al. 2007) and baseline brain activity and metabolism (Wanet-Defalque et al. 1988; Veraart et al. 1990; Jiang et al. 2015) in these regions, and superior attention performance (Roder et al. 1999; Collignon et al. 2006; Forster et al. 2007; Collignon and De Volder 2009) in the blind.

### Functional reorganisation of the attention-related networks and brain maturity

In the blind, the structural and functional reorganisation of the visual cortex has been found to be dependent on the development maturity degree of the visual cortex at the time of onset of blindness. Before or within the critical developmental period, visual deprivation results in more significant reorganisation in the occipital cortex than that occurs after the period. Compared to the LB, the CB/EB have more significant changes in cortical thickness (Jiang et al. 2009; Park et al. 2009; Kupers et al. 2011), glucose metabolism (Wanet-Defalque et al. 1988; Veraart et al. 1990), task-evoked activation (Buchel et al. 1998b), and FCD (Qin et al. 2015) in the occipital cortex. Unexpectedly, we found that the functional reorganisation of the attention-related networks was quite similar among the three blind groups and was not correlated with onset age of blindness, indicating that the functional reorganisation of the attention-related networks is not dependent on their development maturity degree at the time of onset of blindness. That is, the non-visual sensory experience is enough for driving the development of the functional organisation of the attention networks. The increased functional coupling may be related to enhancing or unmasking connections of the attention-related networks (Kupers et al. 2011; Qin and Yu 2013).

In contrast, although the blind people similarly demonstrated increased intrinsic functional coupling between the attention and visual networks, some hubs and connections between the two networks can also impacted by blindness duration. The most vulnerable attention area is the posterior IFG, the conjunction hub of DAN and VAN (Fox et al. 2006), whose functional coupling with the VN was affected by duration of visual deprivation, indicating the experience-dependent factors may reshape the intrinsic functional connectivity.

## Conclusions

In summary, our findings demonstrated that long-term visual deprivation can reshape the intrinsic functional coupling within the attention network, and between the attention and visual networks; furthermore, the functional reorganisation of the attention-related networks is not dependent on their development maturity degree at the time of onset of blindness. These compensatory changes may help blind people to more effectively allocate the remaining sensory resources to be aware of the surrounding environment.

## Acknowledgments

This work was supported by the Natural Science Foundation of China (81771818 and 81425013), National Key Research and Development Program of China (2018YFC1314300), and Tianjin Key Technology R&D Program (17ZXMFSY00090).

