## Supplemental Figures for "Intrinsic functional reorganisation of the attention network in the blind"

**Supplementary Figures**

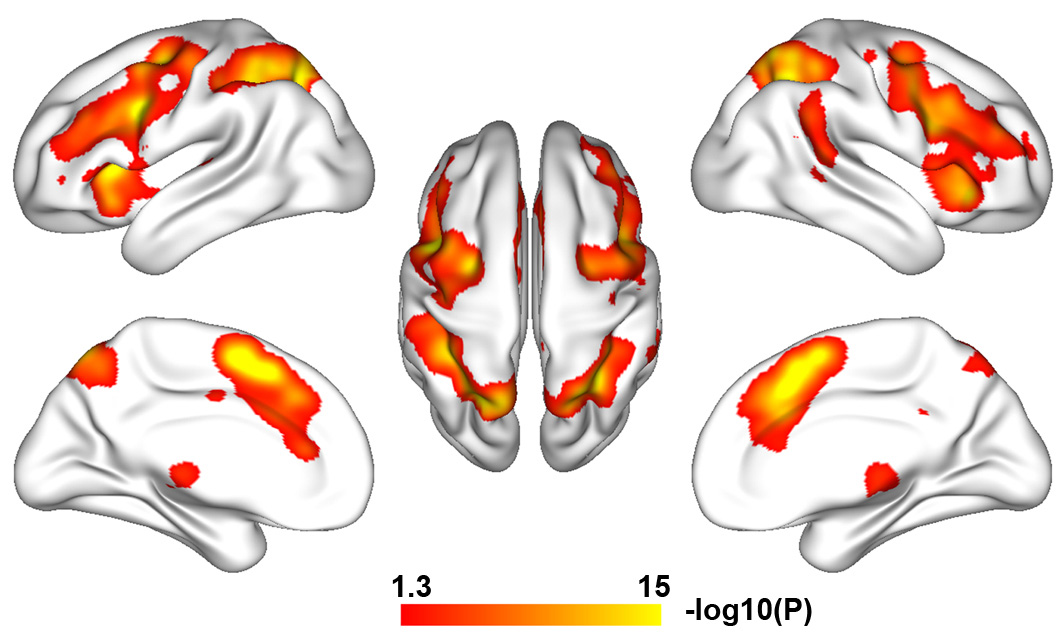

**Figure S1: Spatial distribution of the attention network derived from a meta-analysis of attention-evoked coactivation.** A total of 128 papers that included 360 experiments, 4793 subjects, and 3470 foci were finally included in the meta-analysis based on activation likelihood estimation algorithm (*q* < 0.05, FDR corrected). Color bar represents the logarithmic-transformed *P* value. Abbreviations: FDR = false discovery rate.

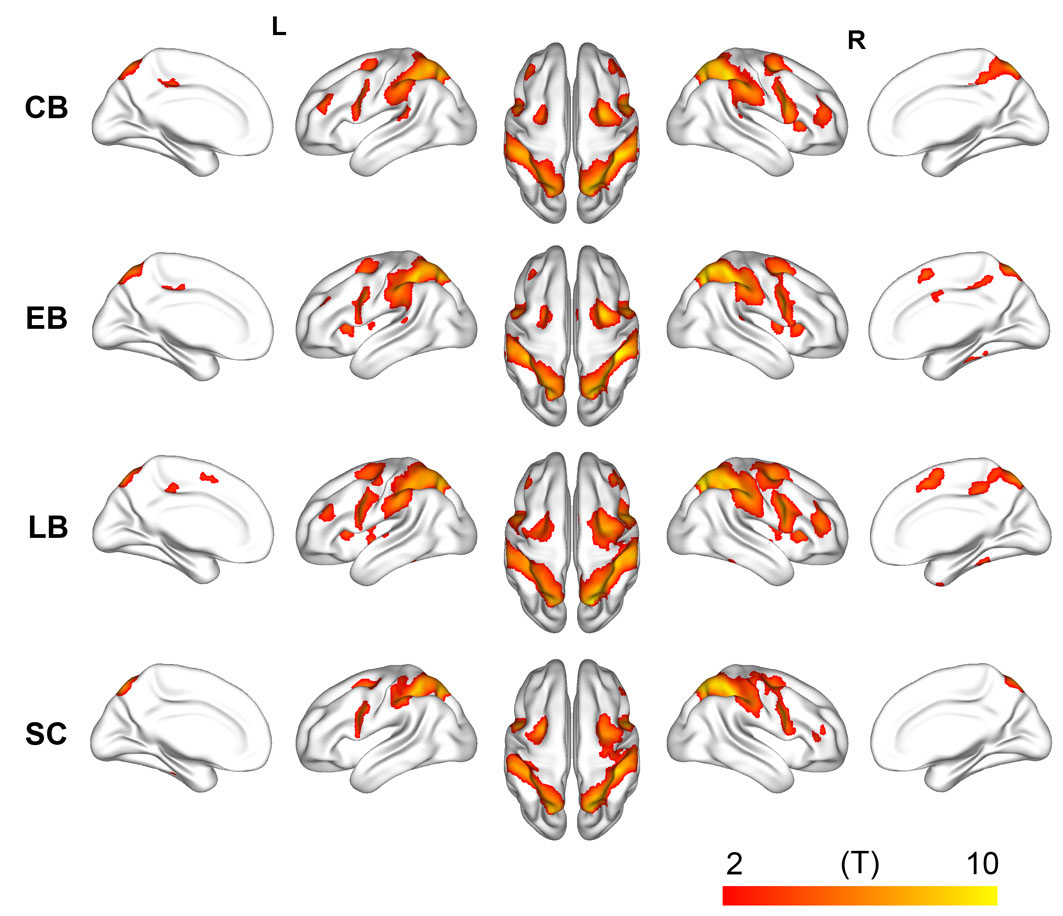

**Figure S2: Dorsal attention network derived from rsFC-based conjunction analysis in each group.** The DAN is constructed by conjunction analyses of the rsFC of two DAN hubs (right FEF and IPS) (*q* < 0.05, FDR corrected). The peak MNI coordinates of the right FEF [28 -6 52] and IPS [30 -58 46] are defined on the ALE coactivation map. The color bar represents the T value of the conjunction analysis. Abbreviations: ALE = activation likelihood estimation, CB = congenitally blind, DAN = dorsal attention network, EB = early blind, FEF = frontal eye field, IPS = inferior parietal sulcus, LB = late blind, MNI = Montreal Neuroimaging Institute, rsFC = resting-state functional connectivity, SC = sighted controls.

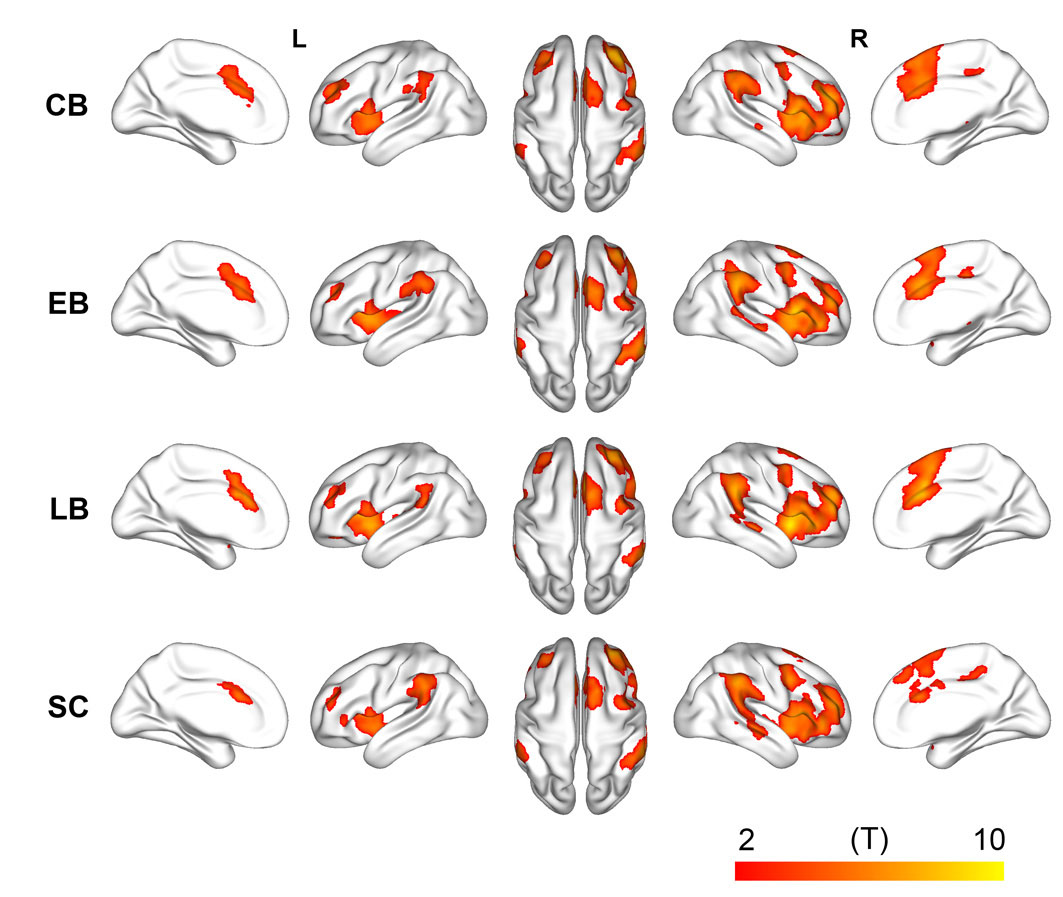

**Figure S3: Ventral attention network** **derived from rsFC-based conjunction analysis in each group.** The VAN is constructed by conjunction analyses of the rsFC of two VAN hubs (right aINS and TPJ) (*q* < 0.05, FDR corrected). The peak MNI coordinates of the right aINS [38 26 -6] and TPJ [58 -42 30] are defined on the ALE coactivation map. The color bar represents the T value of the conjunction analysis. Abbreviations: aINS = anterior insular, ALE = activation likelihood estimation, CB = congenitally blind, EB = early blind, LB = late blind, MNI = Montreal Neuroimaging Institute, rsFC = resting-state functional connectivity, SC = sighted controls, TPJ = temporoparietal junction, VAN = ventral attention network.

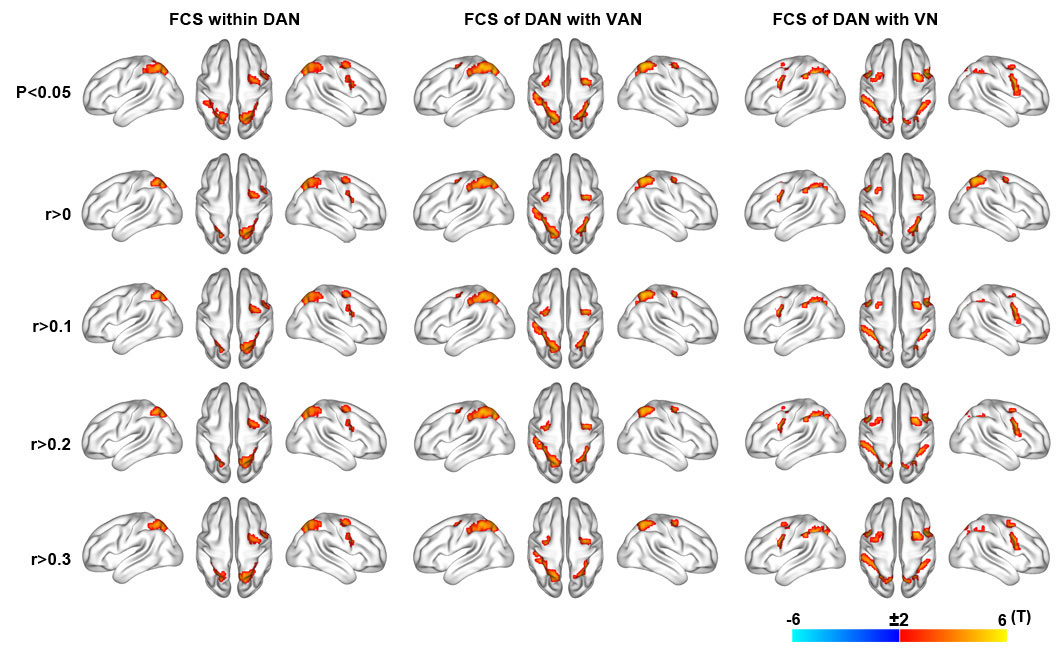

**Figure S4.** **FCS changes of the DAN in the blind with different connectivity thresholds**. General linear model is used to compare FCS differences within a positive rsFC mask between the total blind subjects and the sighted controls (*q* < 0.05, FDR corrected), while controlling for the effects of age, gender, and mean FD. Color bar represents the T value. Warm and cold colors represent increased and decreased FCS in the blind, respectively. The FCS is calculated with different connectivity thresholds (0, 0.1, 0.2 and 0.3). In the text, the FCS is calculated with a connectivity threshold of *P* < 0.05 (uncorrected). Abbreviations: CB = congenital blind, DAN = dorsal attention network, EB = early blind, FCS = functional connectivity strength, FD = frame-wise displacement, FDR = false discovery rate, LB = late blind, rsFC = resting-state functional connectivity, SC = sighted control, VAN = ventral attention network, VN = visual network.

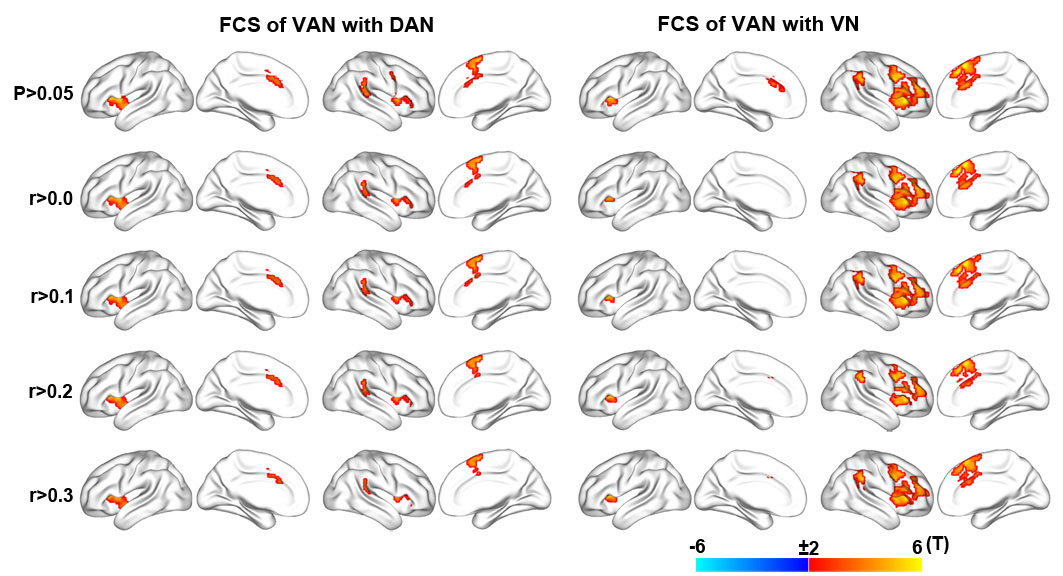

**Figure S5. FCS changes of the VAN in the blind with different connectivity thresholds**. General linear model is used to compare FCS differences within a positive rsFC mask between the total blind subjects and the sighted controls (*q* < 0.05, FDR corrected), while controlling for the effects of age, gender, and mean FD. Color bar represents the T value. Warm and cold colors represent increased and decreased FCS in the blind, respectively. The FCS is calculated with different connectivity thresholds (0, 0.1, 0.2 and 0.3). In the text, the FCS is calculated with a connectivity threshold of *P* < 0.05 (uncorrected). Abbreviations: CB = congenital blind, DAN = dorsal attention network, EB = early blind, FCS = functional connectivity strength, FD = frame-wise displacement, FDR = false discovery rate, LB = late blind, rsFC = resting-state functional connectivity, SC = sighted control, VAN = ventral attention network, VN = visual network.

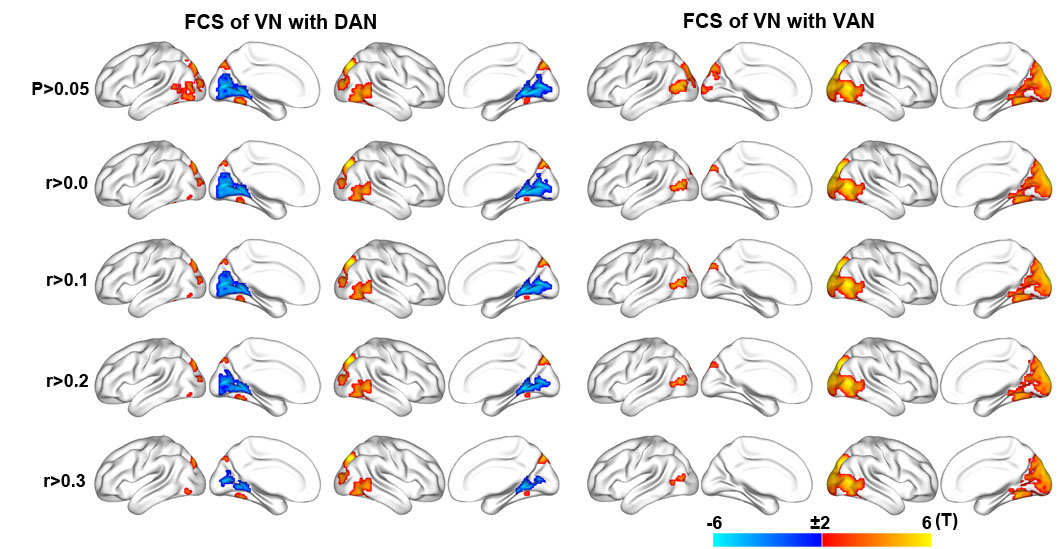

**Figure S6. FCS changes in the VN in the blind with different connectivity thresholds**. General linear model is used to compare FCS differences within a positive rsFC mask between the total blind subjects and the sighted controls (*q* < 0.05, FDR corrected), while controlling for the effects of age, gender, and mean FD. Color bar represents the T value. Warm and cold colors represent increased and decreased FCS in the blind, respectively. The FCS is calculated with different connectivity thresholds (0, 0.1, 0.2 and 0.3). In the text, the FCS is calculated with a connectivity threshold of *P* < 0.05 (uncorrected). Abbreviations: CB = congenital blind, DAN = dorsal attention network, EB = early blind, FCS = functional connectivity strength, FD = frame-wise displacement, FDR = false discovery rate, LB = late blind, rsFC = resting-state functional connectivity, SC = sighted control, VAN = ventral attention network, VN = visual network.

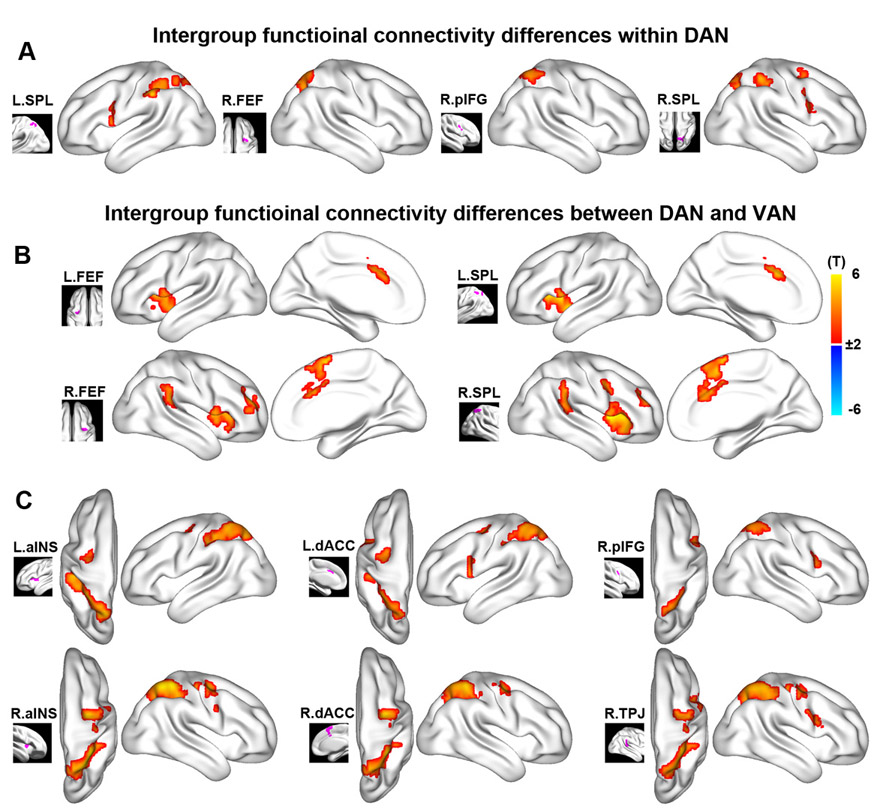

**Figure S7. Specific rsFC changes within the attention networks in the blind.** General linear model is used to compare the seed-based rsFC differences within the attention networks between the total blind subjects and the sighted controls (q < 0.05, FDR corrected), while controlling for the effects of age, gender, and mean FD. Color bar represents the T value. Warm and cold colors represent increased and decreased rsFC in the blind, respectively. The seeds are defined as brain regions with significant changes in FCS within the attention networks in the blind (Fig. 4). (A) shows brain regions within the DAN with significant intergroup differences in rsFC with the seeds of the DAN; (B) shows brain regions within the VAN with significant intergroup differences in rsFC with the seeds of the DAN; and (C) shows brain regions within the DAN with significant intergroup differences in rsFC with the seeds of the VAN. Abbreviations: aINS = anterior insular cortex, CB = congenital blind, dACC = dorsal anterior cingulate cortex, DAN = dorsal attention network, EB = early blind, FCS = functional connectivity strength, FD = frame-wise displacement, FDR = false discovery ratio, FEF = frontal eye field, L, left, LB = late blind, pIFG = posterior inferior frontal gyrus, R, right, rsFC = resting-state functional connectivity, SC = sighted control, SPL = superior parietal lobe, VAN = ventral attention network.

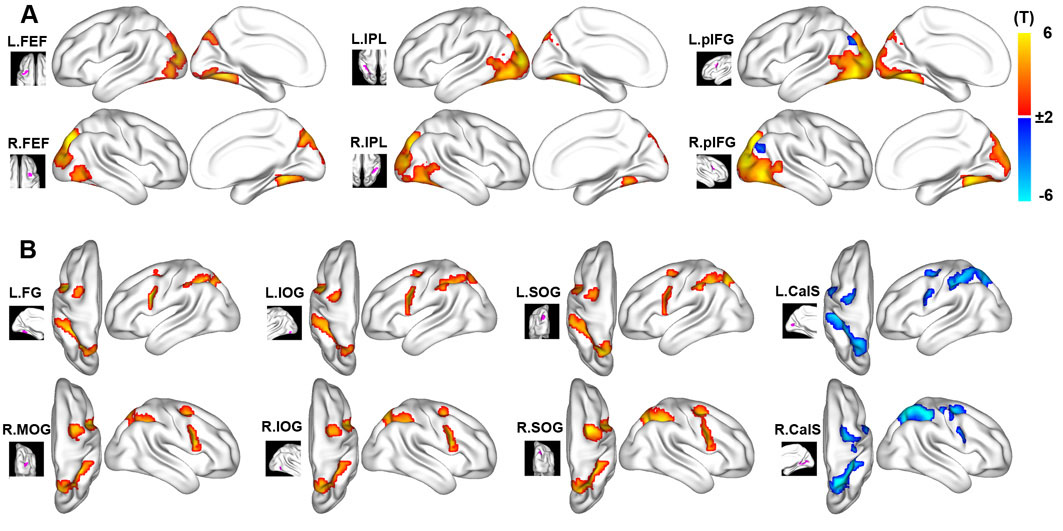

**Figure S8. Specific rsFC changes between the DAN and VN in the blind.** General linear model is used to compare the seed-based rsFC (between the DAN and VN) differences between the total blind subjects and the sighted controls (*q* < 0.05, FDR corrected), while controlling for the effects of age, gender, and mean FD. Color bar represents the T value. Warm and cold colors represent increased and decreased rsFC in the blind, respectively. The seeds are defined as brain regions with significant changes in FCS between the DAN and VN in the blind (Fig. 5). **(A)** shows brain regions within the VN with significant intergroup differences in rsFC with the seeds of the DAN; and **(B)** shows brain regions within the DAN with significant intergroup differences in rsFC with the seeds of the VN. Abbreviations: CalS = calcarine sulcus, CB = congenital blind, DAN = dorsal attention network, EB = early blind, FCS = functional connectivity strength, FD = frame-wise displacement, FDR = false discovery ratio, FEF = frontal eye field, FG = fusiform gyrus, IOG = inferior occipital gyrus, IPL = inferior parietal lobule, L, left, LB = late blind, pIFG = posterior inferior frontal gyrus, R, right; rsFC = resting-state functional connectivity, SC = sighted control, SOG = superior occipital gyrus, VN = visual network.

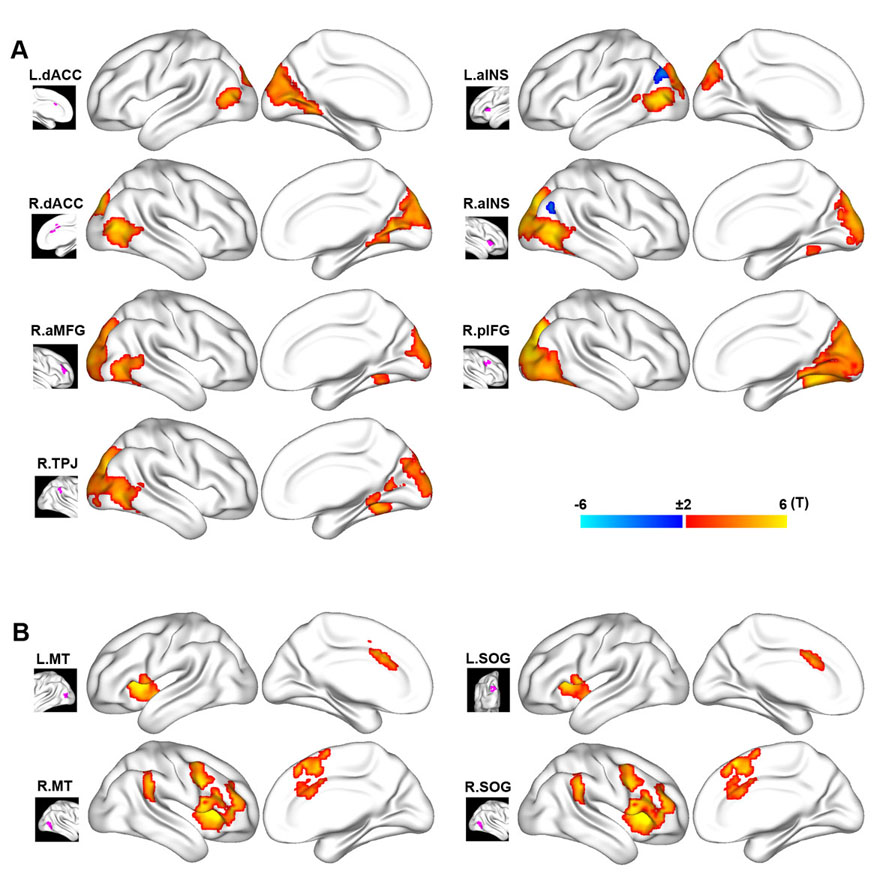

**Figure S9. Specific rsFC changes between the VAN and VN in the blind.** General linear model is used to compare the seed-based rsFC (between the VAN and VN) differences between the total blind subjects and the sighted controls (*q* < 0.05, FDR corrected), while controlling for the effects of age, gender, and mean FD. Color bar represents the T value. Warm and cold colors represent increased and decreased rsFC in the blind, respectively. The seeds are defined as brain regions with significant changes in FCS between the VAN and VN in the blind (Fig. 5). **(A)** shows brain regions within the VN with significant intergroup differences in rsFC with the seeds of the VAN; and **(B)** shows brain regions within the VAN with significant intergroup differences in rsFC with the seeds of the VN. Abbreviations: aINS = anterior insula cortex, aMFG = anterior middle frontal gyrus, CB = congenital blind, dACC = dorsal anterior cingulate cortex, EB = early blind, FCS = functional connectivity strength, FD = frame-wise displacement, FDR = false discovery ratio, L, left, LB = late blind, MT = middle temporal visual area, pIFG = posterior inferior frontal gyrus, R, right; rsFC = resting-state functional connectivity, SC = sighted control, SOG = superior occipital gyrus, TPJ = temporoparietal junction, VAN = dorsal attention network, VN = visual network.
